# Human blastocyst formation in an embryo culture medium mimicking human uterine pH and fluid composition

**DOI:** 10.1101/2024.11.19.623474

**Authors:** Miriam S. Zagers, Maitane Laverde, Majid Tarahomi, Hubert Joris, Sebastiaan Mastenbroek

## Abstract

**Study question:** To study the development of human embryos in a culture medium with pH 6.8 and composition based on previous *in vivo* human uterine measurements.

**Summary answer:** The blastocyst formation rate of donated surplus human preimplantation embryos in embryo culture medium with pH 6.8 and uterine composition was similar to control medium.

**What is known already:** *In vitro* culture conditions can affect human preimplantation embryo development and fertility treatment success rates. The current IVF culture conditions are derived from using animal models, laboratory experience and limited human *in vivo* evidence. Therefore, we previously measured human *in vivo* uterine conditions on the third day following a positive LH test or ovum pick-up. Strikingly, the uterine pH at this time of the menstrual cycle (pH 6.8) appeared lower than the pH used for human preimplantation embryo culture in IVF laboratories worldwide (pH 7.2-7.4). Mimicking *in vivo* conditions may potentially improve *in vitro* embryo development, implantation rates, and overall fertility treatment and safety outcomes.

**Study design, size, duration:** In this preclinical pilot study, 404 donated surplus human day 3 and day 4 embryos were cultured in one of four culture media: ULPC medium (uterine low pH and uterine fluid composition), UC medium (standard pH and uterine fluid composition), ULP medium (uterine pH and standard composition), or control medium (G-2 PLUS medium with standard pH and composition). Blastocyst formation rate on day 5 (after one or two days in culture) was the primary outcome to assess embryo development.

**Participants/materials, setting, methods:** Embryos were randomly allocated to the different culture conditions, stratified by sibling status and maternal age, before thawing. Day 3 and day 4 embryos were separately randomized, ensuring that each culture group contained an equal amount of both day 3 and day 4 embryos. All procedures and culture conditions were similar between groups, except for pH and medium composition. Embryo morphology was assessed immediately after thawing and on developmental day 5 for the primary outcome. Assessors were blinded to medium allocation.

**Main results and the role of chance:** All four culture media supported human preimplantation embryo development into blastocysts. Blastocyst formation rates on day five were 54/102 (52.9%) in ULPC medium, 44/104 (42.3%) in UC medium, 48/98 (49.0%) in ULP medium and 47/102 (46.1%) in control medium.

**Limitations, reasons for caution:** This study is limited by the use of donated surplus human day three and day four embryos. The effect of these new *in vitro* conditions on embryo development during the first three days of human embryo culture is not yet known.

**Wider implications of the findings:** The outcomes of this study indicate the need for further research on mimicking the *in vivo* uterine conditions for *in vitro* embryo culture using the conditions proposed here, including a lower pH (of 6.8) than currently used in IVF laboratories (pH 7.2-7.4).

## Introduction

Worldwide, approximately 1 million babies are born each year as a result of an estimated 4 million embryo transfers (ESHRE, 2023). A critical determinant of success of *in vitro* fertilization (IVF) or intracytoplasmic sperm injection (ICSI) treatments is the quality of the available preimplantation embryos, typically assessed by their morphology. As the embryo development and quality are influenced by the *in vitro* culture environment, ongoing research focuses on enhancing these conditions to potentially increase the success rates of IVF/ICSI. The significance of optimizing the *in vitro* embryo culture environment was highlighted by findings indicating that the embryo culture medium not only impacts the cumulative live birth rate, but also other important IVF outcomes, such as birthweights of children conceived via IVF/ICSI (Dumoulin et al., 2010, Mantikou et al., 2013, Youssef et al., 2015, Kleijkers et al., 2016). Despite ongoing requests for full transparency from the field, manufacturers have consistently refrained from disclosing the exact composition of their embryo culture media (Biggers, 2000, Evers, 2016, Sunde et al., 2016, Tarahomi et al., 2019, Paulson, 2023, Zagers et al., 2023, Paulson and Adashi, 2024). This hinders scientific efforts to understand the effects of the embryo culture medium composition and optimization of this composition for improved human embryo culture. Multiple studies therefore determined at least part of the composition of commercial embryo culture media (Dyrlund et al., 2014, Morbeck et al., 2014a, Morbeck et al., 2014b, Morbeck et al., 2017, Tarahomi, et al., 2019, Zagers, et al., 2023). The observed variability in the compositions of these embryo culture media underscore the absence of a universally acknowledged and evidence-based composition (Zagers, et al., 2023). Concentrations of part of the components appeared to be based on observations from experimental studies with animal models or clinical laboratory experience, while others seemed to mimic concentrations that had been determined in limited samples of *in vivo* oviduct and uterine fluid – before more recent *in vivo* evidence became available (Morbeck, et al., 2014a, Morbeck, et al., 2017, Tarahomi, et al., 2019, Zagers, et al., 2023). A comparison of *in vitro* embryo culture medium compositions with more recent *in vivo* findings revealed that concentrations of some key components in the culture media appear non-physiological, as they differ from the concentrations identified in *in vivo* human oviduct or uterine (Kermack et al., 2015, Utsunomiya et al., 2022, Zagers, et al., 2023, Tarahomi et al., 2024). The discrepancies between *in vitro* human embryo culture conditions and the *in vivo* human embryo environment may be a consequence of the use of animal models (mainly mouse) for culture medium development and optimization (Esfandiari and Gubista, 2020). These findings suggest that more knowledge of the *in vivo* human preimplantation embryo environment may provide leads for optimization of human preimplantation embryo culture medium.

To increase the evidence on the *in vivo* preimplantation embryo environment, we previously measured human uterine temperature and pH on the third day post-LH surge or ovum pick-up and simultaneously aspirated uterine fluid for composition analysis (Tarahomi, et al., 2024). Remarkably, we found the human uterine pH at this time of the menstrual cycle (pH 6.8) to be lower than the pH that is commonly used for human preimplantation embryo culture in IVF laboratories worldwide (pH 7.2-7.4). Our observation seems supported by earlier *in vivo* studies that also reported a lower average pH in the human uterus (pH 6.5-7.1) (Feo, 1955, Sedlis et al., 1967, Yedwab et al., 1976).

The importance of the correct pH for *in vitro* embryo culture and the complexity of determining this correct pH was previously outlined (Swain, 2010, Swain, 2012, Gatimel et al., 2020). The concentrations of most uterine fluid components identified in our *in vivo* measurements aligned with other recent *in vivo* studies (Kermack, et al., 2015, Utsunomiya, et al., 2022). The concentrations of some of the components differed, though, from those in commonly used embryo culture media (Zagers, et al., 2023).

Embryo culture media that best mimic the measured *in vivo* conditions (uterine pH and uterine fluid composition) may potentially improve human preimplantation embryo development and quality, and with that overall fertility treatment and safety outcomes. We developed a human embryo culture medium based on human uterine characteristics that we previously determined (pH 6.8 and the concentrations of uterine fluid components). In the present study we evaluate whether these *in vivo* based conditions support human preimplantation embryo development *in vitro*.

## Materials and Methods

### Embryo culture media

We developed an *in vivo* evidence-based human embryo culture medium (ULPC medium), based on the uterine conditions determined in our previous study in collaboration with Vitrolife, Sweden, (Tarahomi, et al., 2024). We also formulated two additional culture media where only composition (UC medium) or pH (UP medium) differed from control medium. G-2 PLUS was selected as control medium. Thus, four different media were used in this study: ULPC medium with a uterine pH of 6.8 and a uterine fluid composition; UC medium with a standard pH of 7.3 and a uterine fluid composition; ULP medium with a uterine pH of 6.8 and a standard (G2 PLUS) composition; and G-2 PLUS medium maintaining both a standard pH of 7.3 and a standard composition. Vitrolife, Sweden, prepared the three custom made media as follows: UC medium was prepared by emulating the concentrations of 29 uterine fluid components as determined *in vivo*. Some of the 37 human uterine fluid components we measured were not used for the preparation of the ULPC and UC media due to different considerations. Citrulline, ornithine and immunoglobulins were present in human uterine fluid, but excluded from the embryo culture medium for different reasons. Their inclusion could restrict the medium’s future use in patient care due to regulatory requirements. We also decided to not add immunoglobulins to the culture media, as our rationale is that they might have an effect on the endometrium *in vivo*, but we do not expect them to have an effect on embryo culture *in vitro*. Sodium chloride concentration was adjusted to maintain the osmolality below 270 mOsm/kg. This osmolality is in line with the considered optimal range of 260-300 mOsm/kg for *in vitro* human embryo culture. Sodium bicarbonate was used to achieve a standard pH of 7.3 within a 6.0% CO_2_ incubator environment. Cysteine, gentamycin and hyaluronan were also added to the medium. To prepare ULPC medium, HCl was added to half of the volume of UC medium until a pH of 6.8 was achieved. The ULP medium was created by reducing the pH of G-2 PLUS medium to pH 6.8 through the addition of HCl. Human Serum Albumin (HSA) was added to the media as protein source.

### Culture medium quality assessment

Standard culture medium quality assessment was performed by Vitrolife, Sweden, directly after production. This evaluation comprised pH measurement (either pH 7.3 or pH 6.8) and osmolality testing (not exceeding 270 mOSm/kg). Mouse embryo assays (MEAs) were employed to verify the support of mouse embryo development by the culture media. This involved culturing thirty embryos from F1 mouse strains in each lot number of the four embryo culture media, assessing both the proportion of expanded blastocysts and the cell count within each blastocyst after 96 hours.

Complementary analysis of culture medium composition was carried out at Amsterdam UMC, utilizing a Cobas 8000 chemistry analyser (Roche Diagnostics) and ultra-performance liquid chromatography-tandem mass spectrometry (UPLC-MS/MS; Acquity-Quattro Premier XE, Waters). This composition analysis mirrored the method we previously applied to determine human uterine fluid composition (Tarahomi, et al., 2024), confirming that concentrations were similar to those found in human uterine fluid. Additionally, pH levels of each lot number of the four media were determined using an RI pH meter 3 (Cooper Surgical) inside the incubator over four consecutive days. For accurate pH measurement, the meter was calibrated through a three-point calibration process using RI pH 7 and pH 10 buffer solutions (RI calibration maintenance pack 7-90-120; Cooper Surgical), following the manufacturer’s guidelines. To prevent CO_2_ interference with the pH 7.00 buffer solution at 37.0°C during the last step, the cap hole was sealed with tape. Then, with a clean pH probe positioned downwards in the holder, 700 μL of culture medium and 150 μL of mineral oil (Irvine Scientific) were pipetted into the holder, which was then capped, allowing CO_2_ interaction with the bicarbonate in the medium. Within an incubator atmosphere containing 6.0% CO_2_, the pH levels of ULPC and ULP media were measured as 6.8±0.2, and the pH levels of UC and G-2 PLUS media were measured as 7.3±0.2. The pH meter’s error margin is 0.1 pH unit. To account for potential pH drift caused by protein clotting on the pH probe, the pH of the pH 7.00 buffer solution was measured over four days before and after testing four culture media for four days (one lot number of each culture medium).

### Human preimplantation embryos

We used five hundred and seventy-nine surplus human preimplantation embryos, cryopreserved at developmental day 3 (n= 128) or day 4 (n= 451) that were donated for scientific research. Four hundred and fifty-one day 4 embryos were obtained from the IVF laboratory at the Center for Reproductive Medicine of Amsterdam UMC, location University of Amsterdam, in Amsterdam, the Netherlands, cryopreserved between 1999 and 2019. The remaining one hundred and twenty-eight day 3 embryos were obtained from the IVF laboratory at the Center for Reproductive Medicine of Radboudumc in Nijmegen, the Netherlands, cryopreserved in 2013 and 2014.

### Embryo randomization and culture dish preparation

Before thawing, we randomly allocated the embryos to one of the four culture medium groups (ULPC medium, UC medium, ULP medium or control medium) and assigned each embryo a unique study number. We stratified for sibling status and maternal age by simultaneously randomizing sibling embryos using sealed envelopes in blocks of four. Culture dishes were labelled with the study number and prepared with six drops of 25 μL of the allocated embryo culture medium — three drops for washing after thawing, one spare drop, and one drop for medium collection – all covered with 8 mL mineral oil, for overnight equilibration.

### Embryo culture

The next day, we thawed the randomized embryos following the local protocols and transferred each embryo to the corresponding 60×15 mm culture dish containing the allocated medium. The thawed embryos were then cultured in a 33-liter water-jacketed CO_2_/O_2_ triple gas incubator (IKS International), with the incubator settings maintained at 37.0°C, 6.0% CO_2_, and 5.0% O_2_. Throughout the culture period, all culture parameters were kept constant and between the study groups only pH and culture medium composition differed.

### Embryo morphology assessment

Embryos were initially scored immediately post-thawing, the inclusion criteria to accept participation in this study were set at ≥5 blastomeres and ≥50% viability; embryos that did not meet these criteria were directly excluded from the study and discarded. Subsequent embryo assessments were conducted every 24 hours. All day 3 embryos were evaluated after thawing, at day 4 and at day 5. All day 4 embryos were assessed after thawing and at day 5. During each assessment, the number and size of blastomeres and the percentage of fragmentation was recorded for cleavage stage embryos. Morula’s were classified as M1, M2, or M3, and all blastocysts were scored in accordance with the Gardner criteria. Additionally, a video capturing the complete embryo was recorded immediately after morphological assessment, using a z-stack movement to ensure thorough documentation. Two independent embryologists confirmed embryo scores upon the videos.

### Statistical analyses

Logistic regression was used to calculate the odds ratios (OR) and 95% confidence intervals (CI) for blastocyst formation on day 5 in ULPC, ULP, and UC media, with the control medium used as the reference group.

## Results

We developed a new human embryo culture medium, ULPC medium, based on uterine conditions we previously determined in the human uterus (Tarahomi, et al., 2024), and determined whether these uterine characteristics support human embryo development *in vitro*. Table 1 presents the exact composition of ULPC and UC medium, developed to closely mimic the uterine fluid composition as we previously identified (see Materials and Methods for more details). The exact formulation of G-2 PLUS and ULP medium has not been disclosed by the manufacturer, however, we previously determined the concentrations of various components (Zagers, et al., 2023). The UC and G-2 PLUS media had a standard pH of 7.3, and HCl was added to adjust these media to a uterine pH of 6.8, resulting in the preparation of ULPC and ULP media, respectively. The blastocyst formation rate on day 5 was used as the primary outcome measure to assess embryo development in each culture environment.

**Table 1.** The formulations for ULPC and UC medium, based on our previous findings on human uterine fluid composition. HCl was added to this composition to decrease the pH of ULPC medium to pH 6.8.

| Chemical | g/L used to prepare the media with uterine composition |
| --- | --- |
| Sodium chloride | 4,61698120 |
| Potassium chloride | 1,28375100 |
| Sodium dihydrogenphospate 1 hydrate | 0,45536700 |
| Magnesium sulphate heptahydrate | 0,45104010 |
| Calcium chloride dihydrate | 0,22786550 |
| D+ glucose anhydrous | 0,91881600 |
| Lactic acid sodium salt | 0,73959600 |
| Pyruvic acid sodium salt | 0,00880320 |
| Alanyl-glutamine | 0,04921554 |
| L-Alanine | 0,04576792 |
| L-Aspartic acid | 0,03062196 |
| L-Asparagine | 0,00433750 |
| L-Glutamic acid | 0,17094446 |
| L-Glycine | 0,07169124 |
| L-Proline | 0,02542876 |
| L-Serine | 0,02313031 |
| L-Methionine | 0,00570728 |
| Arginine | 0,02374694 |
| Cysteine | 0,00878200 |
| Histidine | 0,01818722 |
| Isoleucine | 0,01289798 |
| Leucine | 0,02532762 |
| Lysine | 0,04360555 |
| Phenylalanine | 0,01658508 |
| Thryptophan | 0,00765454 |
| Threonine | 0,01156767 |
| Tyrosine | 0,01561858 |
| Valine | 0,02222120 |
| Gentamycin | 0,25750000 |
| Sodium bicarbonate | 2,10025000 |
| Albumin | 51,44560000 |
| Hyaluronan | 0,15000000 |

We randomly allocated a total of 579 donated surplus cryopreserved human day 3 (n= 128) and day 4 (n= 451) embryos, stratified for sibling status and maternal age, to one of the four culture medium groups. Day 3 and day 4 embryos were randomized as separate groups, resulting in a balanced distribution of both day 3 and day 4 embryos across the medium groups, accounting for the variability in developmental stages. Four-hundred and four embryos (61 day 3 and 343 day 4 embryos) survived the thawing process, met our inclusion criteria and were included in this study (Table 2a). All included embryos contained 5 or more viable blastomeres or were already in the morula or early blast stadium and showed 50-100% viability (Table 2b). The average maternal age corresponding with all included embryos per culture medium group was: 33.3 years in ULPC medium, 33.7 years in UC medium, 33.5 years in ULP medium and 33.1 years in the control medium. As we randomized the embryos before thawing, equal inclusion of the amount of post-thaw survived embryos and the quality of the embryos after thawing in each culture medium group could not be controlled. To evaluate the distribution of good and lower quality embryos over the culture medium groups, we assessed the embryo quality immediately after thawing. Each culture medium group exhibited 65.3% to 73.5% good quality embryos immediately after thawing on day 3 or 4. Embryos were considered good quality when they contained 8 or more viable blastomeres in combination with a fragmentation score of zero to a maximum of 20%, or when they were scored as morula 1-2 or early blast (Table 2 and Figure 1).

**Table 2.**
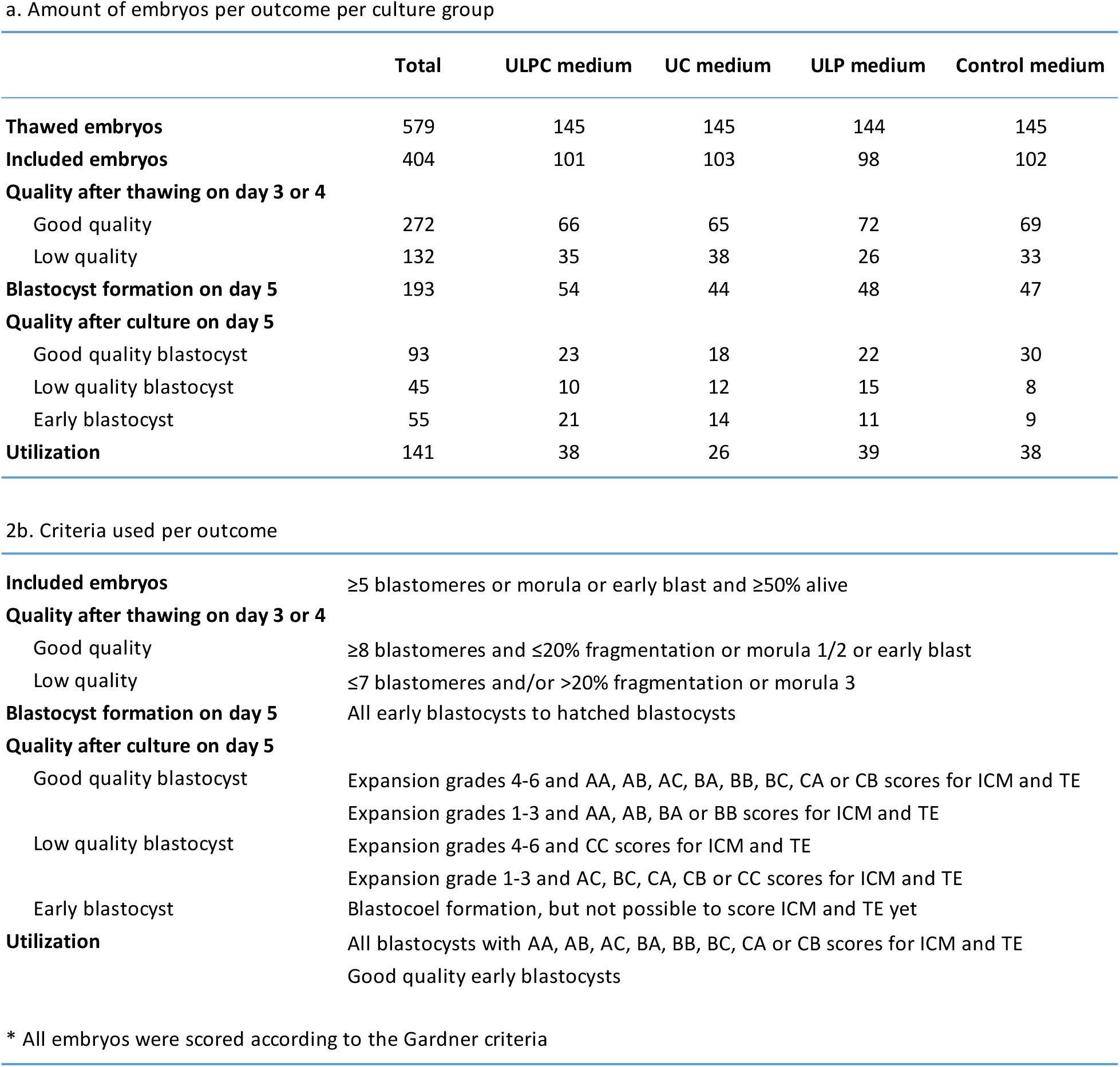
Embryo outcomes. **a**. Amount of embryos per outcome per culture group. **b**. Criteria used per outcome

|  | Total | ULPC medium | UC medium | ULP medium | Control medium |
| --- | --- | --- | --- | --- | --- |
| **Thawed embryos** | 579 | 145 | 145 | 144 | 145 |
| **Included embryos** | 404 | 101 | 103 | 98 | 102 |
| **Quality after thawing on day 3 or 4** |  |  |  |  |  |
| Good quality | 272 | 66 | 65 | 72 | 69 |
| Low quality | 132 | 35 | 38 | 26 | 33 |
| **Blastocyst formation on day 5** | 193 | 54 | 44 | 48 | 47 |
| **Quality after culture on day 5** |  |  |  |  |  |
| Good quality blastocyst | 93 | 23 | 18 | 22 | 30 |
| Low quality blastocyst | 45 | 10 | 12 | 15 | 8 |
| Early blastocyst | 55 | 21 | 14 | 11 | 9 |
| **Utilization** | 141 | 38 | 26 | 39 | 38 |

2b. Criteria used per outcome
|  |  |
| --- | --- |
| **Included embryos** | ≥5 blastomeres or morula or early blast and ≥50% alive |
| **Quality after thawing on day 3 or 4** |  |
| Good quality | ≥8 blastomeres and ≤20% fragmentation or morula 1/2 or early blast |
| Low quality | ≤7 blastomeres and/or >20% fragmentation or morula 3 |
| **Blastocyst formation on day 5** | All early blastocysts to hatched blastocysts |
| **Quality after culture on day 5** |  |
| Good quality blastocyst | Expansion grades 4-6 and AA, AB, AC, BA, BB, BC, CA or CB scores for ICM and TE<br>Expansion grades 1-3 and AA, AB, BA or BB scores for ICM and TE |
| Low quality blastocyst | Expansion grades 4-6 and CC scores for ICM and TE<br>Expansion grade 1-3 and AC, BC, CA, CB or CC scores for ICM and TE |
| Early blastocyst | Blastocoel formation, but not possible to score ICM and TE yet |
| **Utilization** | All blastocysts with AA, AB, AC, BA, BB, BC, CA or CB scores for ICM and TE<br>Good quality early blastocysts |

\* All embryos were scored according to the Gardner criteria

**Figure 1.**
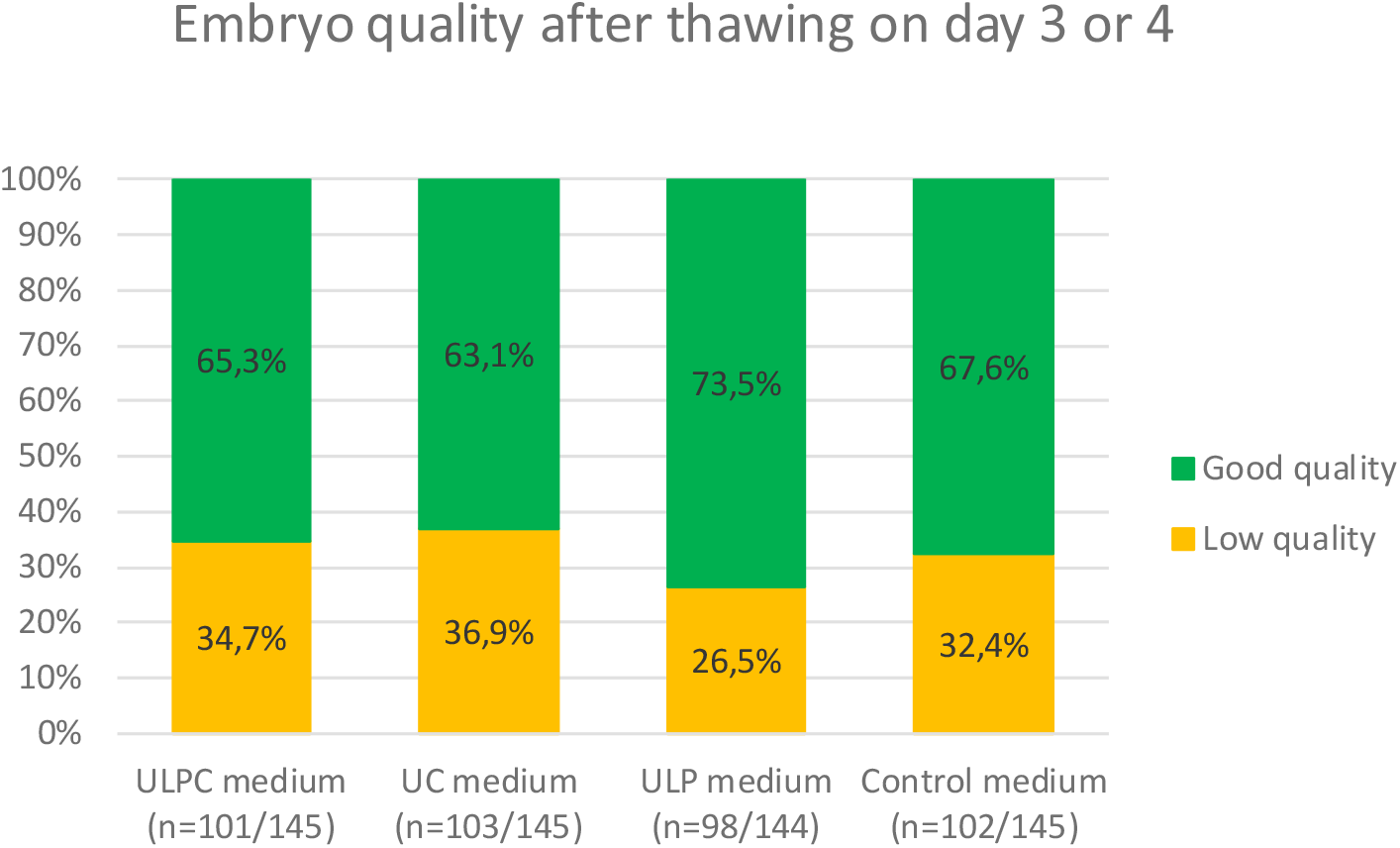
Embryo quality of all included embryos after thawing on developmental day 3 or 4 per culture group. The quality criteria are described in table 1b. ULPC medium: uterine pH 6.8 and uterine fluid composition; UC medium: standard pH 7.3 and uterine fluid composition; ULP medium: uterine pH 6.8 and standard G-2 PLUS composition; Control medium: G-2 PLUS medium with standard pH 7.3 and standard composition.

In total, 193 embryos (47.8%) of the 404 included embryos developed into a blastocyst on day 5 (Table 2). The blastocyst formation rate on day 5 per culture medium group was: 54/101 (53.5%) in ULPC medium, 44/103 (42.7%) in UC medium, 48/98 (49.0%) in ULP medium and 47/102 (46.1%) in the control medium (Table 2a and Figure 2). We used the Gardner criteria for morphological scoring of the blastocysts and determined whether the blastocysts were of good quality or low quality, or still in the early blastocyst stadium, when it is not possible yet to score the inner cell mass and trophectoderm (Figure 2). The criteria we used to categorize the blastocysts are described in Table 2b.

**Figure 2.**
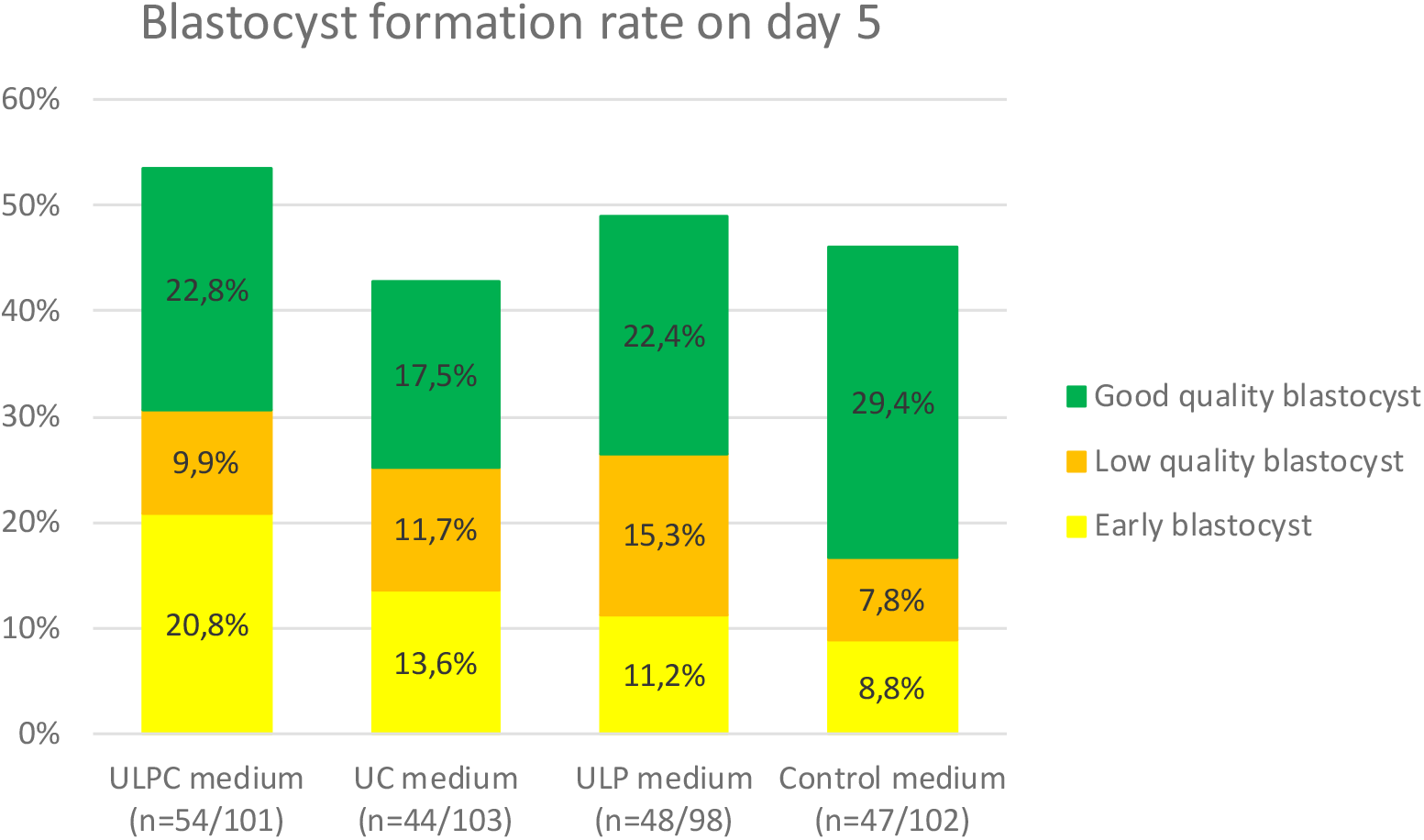
Blastocyst formation rate and blastocyst quality after culture on developmental day 5 per culture group. The quality criteria are described in table 2b. ULPC medium: uterine pH 6.8 and uterine fluid composition; UC medium: standard pH 7.3 and uterine fluid composition; ULP medium: uterine pH 6.8 and standard G-2 PLUS composition; Control medium: G-2 PLUS medium with standard pH 7.3 and standard composition.

The blastocyst formation rate on day 5 was highest in ULPC medium (Figure 2). Most blastocysts were of good quality, however, compared to the control medium group a higher percentage of blastocysts was still an early blastocyst on day 5. The control medium group had the highest percentage of good quality blastocysts. The two intermediate culture media showed different results (Figure 2). Although the blastocyst formation rate on day 5 in ULP medium was similar to ULPC and the control medium, the proportion of blastocysts of lower quality was relatively high in this culture medium group. The blastocyst formation rate on day 5 in UC medium was lower than in the other three culture medium groups. The odds ratios per culture medium compared to the control medium were: ULPC medium OR 1.34 (95% CI 0.77 – 2.33), UC medium OR 0.87 (95% CI 0.50 – 1.52), ULP medium OR 1.12 (95% CI 0.65 – 1.96), highlighting that the chance of blastocyst formation is slightly higher in ULPC and ULP medium and slightly lower in UC medium compared to the control medium. While none of the differences is statistically significant, numbers may be relevant for clinical practice.

We also determined the hypothetical “embryo utilization rate” (EUR) on day 5 (embryos to transfer to the uterus or to cryopreserve for a following cycle) for each medium group: 38/101 (37.6%) in ULPC medium, 26/103 (25.2%) in UC medium, 39/98 (39.8%) in ULP medium and 38/102 (37.3%) in the control medium. The EUR criteria we used are described in Table 2b.

## Discussion

In this study, we tested a new developed *in vivo* evidence-based human embryo culture medium, ULPC medium, that closely mimics our previous findings on the *in vivo* environment of human preimplantation embryos. We showed that donated surplus human day 3 or day 4 embryos cultured in ULPC medium developed into blastocysts and that the blastocyst formation rate on day 5 in this medium was comparable to the blastocyst formation rate on day 5 in the control medium, G-2 PLUS, a common embryo culture medium currently used in IVF laboratories.

We used a back to nature approach, as it can lead to true optimization of human embryo culture media. Although true optimization through fully mimicking *in vivo* conditions remains a long-term goal, since we are currently limited to relying on *in vivo* determined components and constrained by regulatory requirements, this approach holds promise. The resulting ULPC medium is characterized by the integration of *in vivo* findings on uterine fluid composition and uterine pH. Another research group used a similar approach and developed OVIT, a human embryo culture medium based on the analysis of 31 components in human oviduct fluid (Utsunomiya, et al., 2022). In the present study, we assessed for the first time human preimplantation embryo development in a culture medium with pH 6.8, based on the pH we found *in vivo*, and significantly lower than the standard pH of 7.3 used in IVF laboratories worldwide.

An advantage of this study is the use of donated surplus human preimplantation embryos for the assessment of embryo development, instead of using mouse embryos that are commonly used for embryo culture medium development and validation. The number of human preimplantation embryos used here is large enough to ensure that the mimicked uterine conditions support human preimplantation embryo development into blastocysts.

The use of donated surplus IVF embryos is, however, still not ideal and several limitations are associated with the use of these embryos. They were cultured in a different culture environment for the first couple of days of their development, using culture media and conditions that were routinely used in the IVF laboratories between 1999 and 2019, before they were cryopreserved and later donated for research. As the donated surplus embryos were already developed until day 3 or 4 before cryopreservation, the effect of the *in vivo* mimicked conditions on earlier stages of *in vitro* embryo culture is yet to be determined. The embryos used for this study are in general of lower quality than embryos regularly transferred in IVF, as the best quality sibling embryos are commonly used for embryo transfer(s). Consequently, the total post-thaw survival rate (69.8%) was lower than normally reported (74% - 90%) (Edgar and Gook, 2012, Rienzi et al., 2017). However, it needs to be taken into account that the post-thaw survival rate after slow freezing – which was used for these embryos – often results in lower post-thaw survival rates than after vitrification, which is reported to be around 90%. Since most of our previous human uterine measurements were performed on the third day following a positive LH test in a natural menstrual cycle, and an embryo would be expected to be at cleavage stage at that time, one could speculate that the new medium might be favourable for preimplantation cleavage stage development rather than day 4 to day 6 development. However, we also performed eight uterine measurements on the third day following ovum pick-up in women that did not receive a transfer because of potential ovarian hyperstimulation syndrome (OHSS) or elective freeze-all, and found that the uterine pH and composition appeared comparable to the other measurements, undermining that theory (Tarahomi, et al., 2024).

Since it is not allowed by law in the Netherlands to create embryos for research – which would have allowed to study also the first days of embryo development, and would theoretically have resulted in a cohort of embryos of similar quality compared to routine care – we still consider the use of donated surplus embryos the best available model for this study.

The blastocyst formation rate was highest in the ULPC medium group, confirming that this new medium effectively supports human preimplantation embryo development to the blastocyst stage. However, the proportion of good quality blastocysts in this culture medium group appeared to be lower compared to the control group, and a higher percentage of early blastocysts was observed, suggesting that embryo development might progress at a slower rate in the ULPC medium.

The two intermediate media were less effective in supporting human embryo development compared to the ULPC medium. Although ULPC and UC media differed only in the pH of the culture medium, the UC group demonstrated a lower blastocyst formation rate on day 5. This difference cannot be attributed solely to the slightly higher proportion of low-quality embryos immediately after thawing in the UC medium group. It might be that the combination of the uterine fluid composition with a higher pH (pH 7.3) than found *in vivo* (pH 6.8) hampers embryo development in UC medium, resulting in less blastocyst formation. Contrastingly, a lower *in vivo* pH of 6.8 in combination with a G-2 PLUS medium composition in ULP medium did support embryo development into a blastocyst, as the blastocyst formation rate on day 5 in ULP medium was similar to (and not significantly different from) ULPC medium and the control medium. At the same time, the higher proportion of lower quality blastocysts on day 5 in ULP medium might indicate that the combination with the control medium composition is not optimal either. These differences between ULPC medium and the two intermediate media – with only uterine composition or uterine pH – suggest that a combination of a uterine pH with a uterine fluid composition best supports embryo development.

We hypothesize that pH variations might have influenced embryo development and blastocyst quality *in vitro* in the embryo culture media with a lower uterine pH (ULPC medium and ULP medium). As the included embryos were initially cultured in a culture medium with a standard of pH 7.3 before cryopreservation and eventually thawed in a thawing medium buffered to maintain a pH of 7.3, they likely experienced a pH shock upon transfer to a culture dish with either ULPC medium or ULP medium.

This pH shock, due to the sudden environmental pH change when transferring the embryos from the thawing medium to their allocated culture dish with low pH ULPC or ULP medium, might have resulted in slower embryo development or lower blastocyst quality in the ULPC and ULP medium groups. An *in vitro* culture system mimicking the *in vivo* environment from fertilization until embryo transfer to the uterus, perhaps with a gradual decrease in pH as has been discussed before (Swain, 2012), could support embryo development without a pH shock, potentially providing better results. Moreover, IVF/ICSI embryos cultured in a standard pH of 7.3 may experience a similar pH shock after transfer to the uterus. Therefore, the utilizable embryos in the control group, including the large proportion of high quality blastocysts, may have been experienced a similar pH shock in the uterus if they had been transferred to a uterus to result in a pregnancy. At the same time, the embryos are subject to other stressors than pH shock, for example as a consequence of cryopreservation and thawing. It is unclear how the effect of pH shock relates to these other stressors.

Attempting to perfectly mimic the dynamic *in vivo* environment of preimplantation embryos *in vitro* is challenging due to variations in oviduct and uterine fluid compositions among women and during the menstrual cycle. Where we currently only know part of the *in vivo* oviduct and uterine conditions, research into these conditions should be used to enhance our understanding of the natural human preimplantation embryo environment and the crosstalk between a preimplantation embryo and it’s *in vivo* environment until implantation. Given that the natural environment could be considered optimal, the oviduct and uterine pH, fluid composition and other parameters should serve as key factors in guiding the optimization of *in vitro* human embryo culture systems.

## Conclusion

Our results suggest that embryo culture media designed to mimic *in vivo* uterine characteristics, including a lower pH than commonly used in routine IVF practice worldwide, support *in vitro* development of human embryos into blastocysts when tested on donated surplus IVF embryos. This warrants further research that should focus on the complete culture period, from fertilization to embryo transfer. Such research should also examine the effect on IVF/ICSI outcomes, including embryo development, cumulative live birth rates, and the health of the children born through IVF. Additionally, continued investigation of the *in vivo* human preimplantation embryo environment, covering both oviductal and uterine conditions, is still crucial to expand our biological knowledge and to improve our ability to mimic *in vivo* conditions more accurately *in vitro*.

## Data availability

The data underlying this article will be shared on reasonable request to the corresponding author.

## Authors’ roles

M.Z. and S.M. designed the study. M.Z. and M.L. collected all data. M.Z., M.L. and S.M. analysed and interpreted the data. M.Z. wrote the final manuscript. M.Z., M.L., H.J. and S.M. contributed to valuable discussions and provided critical insights over the course of the study. All authors revised the manuscript and approved publication of the last version.

## Acknowledgements

We would like to extend our sincere gratitude to the following individuals for their invaluable contributions to this research project. We are grateful to Dr Annemieke de Melker, Dr Liliana Ramos, and Ing. Emma Brinkmann for their efforts in providing the donated surplus human preimplantation embryos. Special thanks go to Dr Liliana Ramos for also critically reviewing the manuscript prior to publication. We would like to acknowledge Dr Madelon van Wely for her assistance with the statistical analysis. Finally, we express our appreciation to Dr Frédéric Vaz and the Core Facility Metabolomics of the Amsterdam UMC (www.cfmetabolomics.nl), and Dr Femke Schrauwen, Ing. Johannes J. de Groot and the Laboratory General Clinical Chemistry for the measurements of some the components of the embryo culture media used in this study.

## Conflict of interest

We have developed a new culture medium as part of this study, without any financial interest or intent for commercial gain. This commitment is underscored by the decision to publicly share the recipe of the culture medium, ensuring transparency and accessibility for the broader scientific community.

The authors also declare that Vitrolife, Sweden, participated in the development and production of the three embryo culture media mimicking uterine pH and/or uterine fluid composition used in this study. Vitrolife, Sweden, provided all culture media used in this study in-kind. Vitrolife, Sweden, had no role in the study design, data collection, analysis or interpretation of the results.

